# Structure and assembly of CAV1 8S complexes revealed by single particle electron microscopy

**DOI:** 10.1101/2020.06.04.133678

**Authors:** Bing Han, Jason C. Porta, Jessica L. Hanks, Yelena Peskova, Elad Binshtein, Kelly Dryden, Derek P. Claxton, Hassane S. Mchaourab, Erkan Karakas, Melanie D. Ohi, Anne K. Kenworthy

## Abstract

Highly stable oligomeric complexes of the monotopic membrane protein caveolin serve as fundamental building blocks of caveolae. Current evidence suggests these complexes are disc shaped, but the details of their structural organization and how they assemble are poorly understood. Here, we address these questions using single particle electron microscopy of negatively stained recombinant 8S complexes of human Caveolin-1. We show that 8S complexes are toroidal structures ~15 nm in diameter that consist of an outer ring, an inner ring, and central protruding stalk. Moreover, we map the position of the N- and C-termini and determine their role in complex assembly, and visualize the 8S complexes in heterologous caveolae. Our findings provide critical insights into the structural features of 8S complexes and allow us to propose a new model for how these highly stable membrane-embedded complexes are generated.

## Introduction

As early as the 1950s the presence of flask-shaped plasma membrane invaginations was noted in endothelial cells. It is now recognized that the surface of a wide range of cell types are decorated by these 50-100 nm structures known as caveolae (*1*). Caveolae have several unique features that distinguish them from other types of coated vesicles. The membranes in caveolae are not continuously curved, but rather exhibit a polygonal-like appearance (*2, 3*). They are relatively uniform in size and assemble in an all-or-none fashion (*1, 4*). Although they can undergo endocytosis, they spend most of their lifecycle attached to the plasma membrane in the form of open flasks (*1*). They can also undergo reversible flattening to serve as membrane reservoirs (*1, 3, 5*).

The 21 kDa integral membrane protein Caveolin 1 (Cav1) is an essential structural component of caveolae as well as the founding member of the caveolin gene family (*1*). Caveolin family members share a similar membrane topology and contain multiple domains. For the case of Cav1, this includes an N-terminal region (residues 1-101), a hairpin-like intramembrane domain (residues 102-134), and a C-terminal domain (residues 135-178). Contained within the N-terminal region are a predicted unstructured region (residues 1-49) and several partially overlapping domains including a highly conserved signature motif (residues 68-75), an oligomerization domain (residues 61-101) and a scaffolding domain (residues 82-101). The C-terminal domain is thought to fold into a long alpha helix and contains three palmitoylation sites (*6, 7*). It has also been predicted to potentially contain a beta strand in the extreme C-terminal region (*8*). The 33-residue intramembrane domain forms an unusual reentrant helix-break-helix structure (*9*), defining caveolin as a member of the poorly understood class of monotopic membrane proteins (*10*). This hairpin-like domain inserts into the cytoplasmic face of the plasma membrane, placing both the N- and C-terminus in the cytoplasm.

Shortly after Cav1 was first identified as a key component of caveolae, the protein was shown to self-assemble into highly stable high molecular weight oligomers (*11, 12*). These oligomers assemble cooperatively shortly after Cav1 is synthesized (*12*), are estimated to contain 7-14 copies of Cav1(*11–13*), and are ~8S in size when analyzed by velocity gradient centrifugation (*14*). The formation of these oligomers is an essential step in caveolae biogenesis, as mutant versions of the protein that fail to generate oligomers are incapable of supporting caveolae assembly (*15*). When they traffic through the Golgi complex, 8S complexes further oligomerize in a cholesterol-dependent manner into higher-order 70S complexes (*14*). 70S complexes later become incorporated into the plasma membrane where they recruit caveolae accessory proteins such as the cavins that assist in sculpting caveolar membranes, as well as other proteins that regulate caveolae dynamics to ultimately generate mature caveolae (*14, 16*).

Some effort has been made to characterize the structural features of Cav1 monomers, but a three dimensional structure of the protein is still lacking (*6, 7*). Furthermore, little is known about the structure of 8S oligomers or how caveolin monomers are packaged within them. Early studies suggested Cav1 oligomers are organized either into 4-6 nm spherical particles (*11*) or filaments composed of rings (*13*). However, more recent studies point to a disc-like organization of the structures. 8S complexes purified from mammalian cells expressing Cav1 and Cav2 are disc shaped structures 15-17 nm in diameter with a thickness of 5 nm (*2*). The most detailed structural insights into the 8S complexes come from EM analysis of complexes formed by the muscle-specific caveolin family member, Caveolin-3 (Cav3). Cav3 is ~65% identical and ~85% similar to Cav1, and like Cav1, Cav3 assembles into high molecular weight oligomers and drives caveolae formation (*17*). Single molecule EM analysis of negatively stained Cav3 complexes at a resolution of ~17 Å revealed the protein assembles into toroidal structures with a diameter of ~165 Å and a height of ~ 55 Å, and consist of nine copies of Cav3 organized into a wedge-like structure (*18*). Based on gold labeling studies, the C-terminal domains of the monomers have been proposed to be positioned within the center of the complex (*18*).

Together with structural data for the cavins, recent EM structures of Cav1 and Cav3 complexes have led to the current model of caveolae organization (*2, 3*). According to this model, 8S complexes form flat faces of the polyhedral-like membranes of caveolae, and are surrounded by a network of filaments generated by cavins which enforce the polyhedral architecture of caveolar membranes (*2, 3*). Yet key questions about the structure of caveolae and its fundamental building blocks remain unanswered, including how caveolin monomers are packaged within 8S complexes, how the complexes are assembled, whether the overall structural elements of 8S complexes are conserved between caveolin family members, and how they interact with one another and accessory proteins to bend membranes and generate caveolae.

Cav1 is not only capable of inducing caveolae biogenesis in eukaryotes; the expression of mammalian Cav1 drives the formation of caveolae-like structures in *Escherichia coli (E.coli)* (*8, 19*). Termed heterologous caveolae (*h*-caveolae), these caveolin-induced vesicles bud from the inner membrane into the bacterial cytoplasm where they accumulate in the form of closed vesicles. Despite their bacterial origins, *h*-caveolae recapitulate several key properties of mammalian caveolae: they are similar in size, are thought to contain similar numbers of Cav1 molecules, and are composed of high molecular weight Cav1 oligomers (*19*). Furthermore, *h*-caveolae exclude detectable levels of bacterial proteins (*19*). These findings suggest *E. coli h*-caveolae represent a useful model to investigate the structure of Cav1 in a system where it both oligomerizes and induces membrane curvature. Here, we utilized *E. coli* as a model system to study the organization of Cav1 8S complexes, mapped the positions of key regions of the protein, determined which domains contribute to complex assembly, and visualized the 8S complexes within *h*-caveolae.

## Results

### CAV1 forms 8S-like complexes in *Escherichia coli*

Previous work has shown that Cav1 forms high molecular weight complexes when expressed in *E. coli* (*19*), but the molecular organization of these complexes was not determined. To address this question, total membrane proteins were extracted using 2% n-Dodecyl-beta-Maltoside (C12M) from *E. coli* cells expressing wild type human CAV1α with a C-terminal 6X-Histidine tag (WT CAV1). Detergent-solubilized WT CAV1 oligomers were separated from the protein lysate by Ni-sepharose affinity followed gel filtration chromatography. Three major peaks were observed in the elution profile (Figure 1A). SDS-PAGE and Coomassie staining of individual fractions indicate each peak contains a ~22 kDa protein, which is consistent with the molecular weight of CAV1 (Figure 1B). The presence of CAV1 was confirmed by Western blotting of the SDS-PAGE gels (Figure 1C).

**Figure 1.**
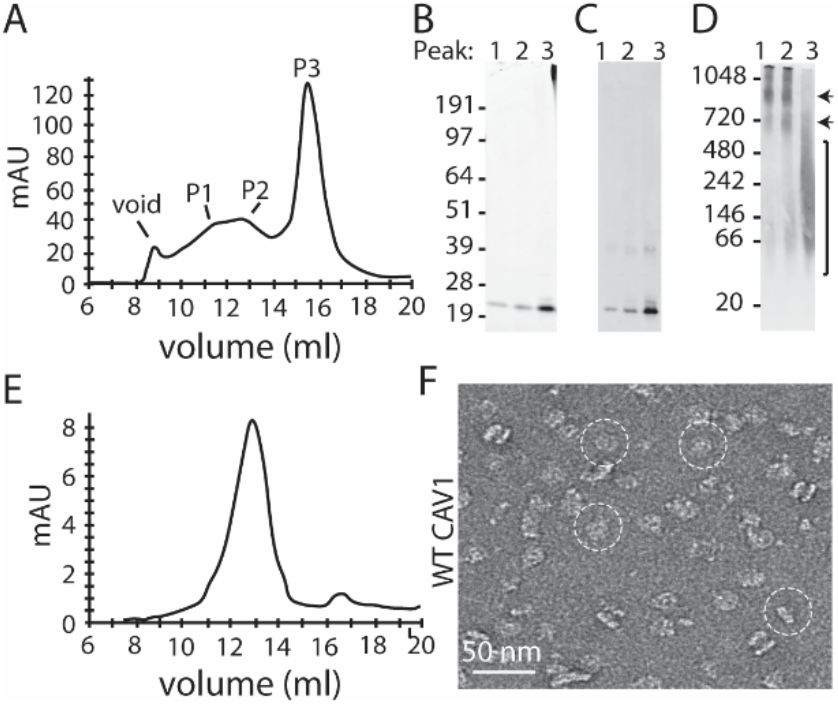
Purification of 8S WT CAV1 complexes from *E. coli*. (**A**) Gel filtration profile of detergent solubilized WT CAV1 oligomers. The position of the void and Peaks P1, P2, and P3 are labeled. (**B**) Coomassie stained SDS-PAGE gel and (**C**) Western blot analysis show that each peak is composed of CAV1. (**D**) Native gel of P1, P2, and P3 shows CAV1 is present in high molecular weight oligomeric complexes. Arrows show location of high molecular weight oligomers typical of 8S-like complexes. Bracket shows location of lower molecular weight oligomeric complexes. (**E**) Gel filtration profile of WT CAV1 oligomers from P2 run a second time over the gel filtration column. One major peak was observed. (**F**) Representative negative stain image of particles from the WT CAV1 peak isolated in E. A few examples of the 8S particles are circled. Scale bar, 50 nm.

A characteristic feature of 8S complexes is their migration as high molecular weight complexes on blue native gels (*15, 20*). To assess the size of the complexes, we performed blue native electrophoresis-based western blot analysis. CAV1 in peaks 1 and 2 (P1 and P2) migrated in multiple high molecular weight complexes with molecular masses greater than 720 kDa in size (Figure 1D), similar to the molecular mass of CAV1 8S complexes extracted from mammalian cells (*20*). Cav1 found in the third peak migrated in lower molecular mass complexes ranging between 40-500 kDa (Figure 1D). These findings suggest CAV1 derived from *E. coli* membranes can generate 8S-like complexes as well as smaller oligomers.

Fractions from the three peaks were further examined using negative stain electron microscopy. All three peaks contained disc-shaped particles (Figure S1). However, the particles in peaks 1 and 2 were larger and appeared more homogenous than the smaller, more heterogenous complexes isolated in Peak 3 (Figure S1).

Further inspection revealed that particles in Peak 1 and 2 are disc shaped. Both “*en face*” and side views of the complexes were apparent, including both individual complexes and dimerized discs (Figure S1A, B). To improve homogeneity of the discs, fractions encompassing Peak 2 were run a second time over gel filtration. One peak was recovered and visualized using negative stain EM (Figure 1E-F). This fraction was used for further analysis of the 8S WT CAV1 complexes.

### CAV1 assembles into disc-shaped complexes that contain a central stalk

To better characterize the structural features of the 8S WT CAV1 complexes, we collected a dataset of ~16,000 negatively stained particles of CAV1. The particles were classified into twenty 2D class averages using RELION (*21*). This analysis revealed classes corresponding to both the *en face* and side views of 8S CAV1 complex (Figure 2A). *The en face* view consists of disc-shaped complexes that are ~15 nm in diameter (Figure S2A and Table S1). The disc structure is composed of a ~4.2 nm wide outer ring and a ~2 nm wide inner ring with strong densities connected by a thinner ring with a weaker density (Figure 2A-B). The side view reveals a T-shaped structure consisting of a thin, flat cap that is ~15 nm wide and ~4.8 nm thick and a central small protruding stalk that is ~3.6 nm wide (Figure 2 A-B). The stalk emanates from the inner ring seen in the *en face* view. Some side view classes showed dimerized 8S CAV1 discs that appeared to be interacting through the stalk region (Figure 2A, bottom right panel).

**Figure 2.**
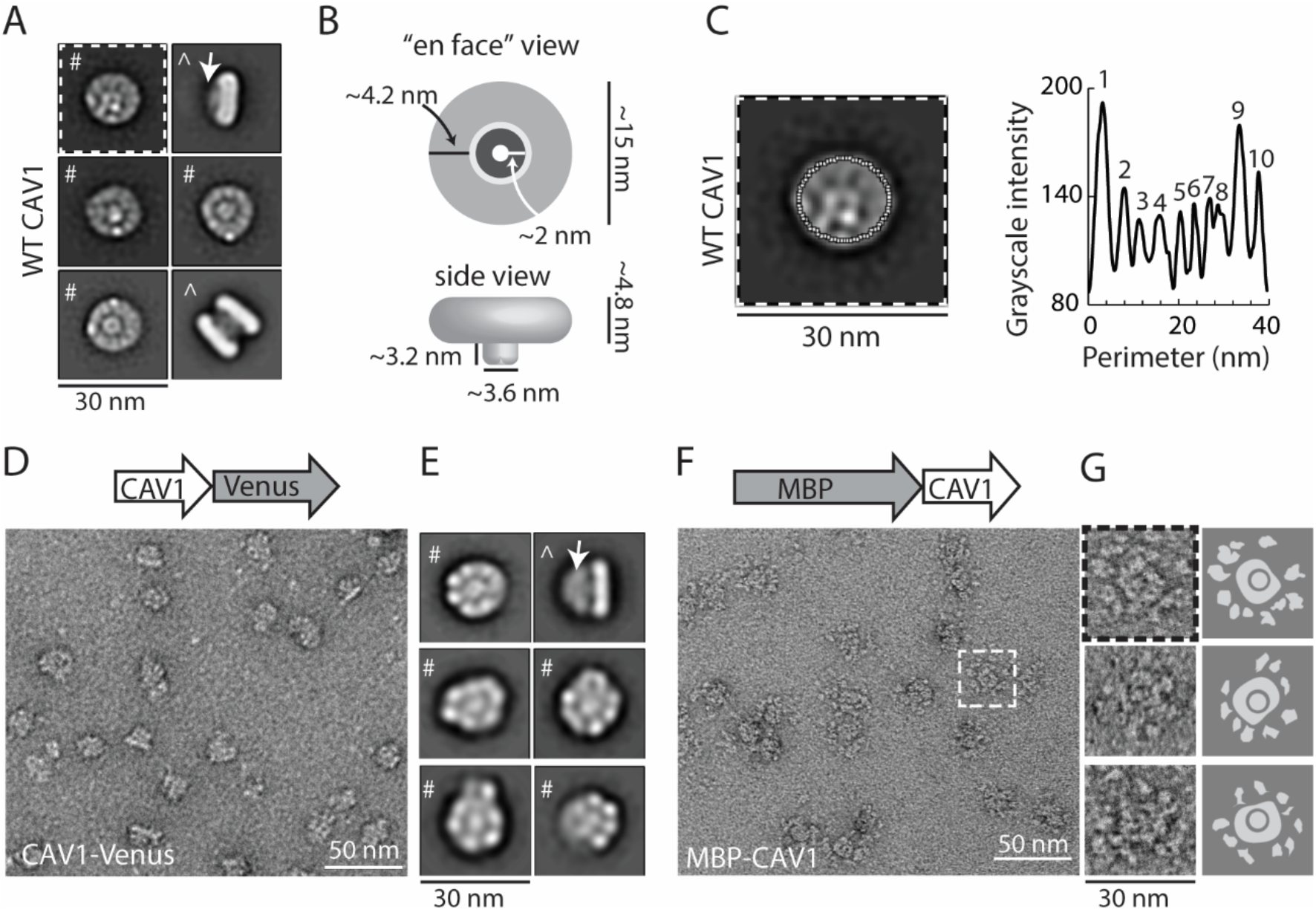
Organization of 8S WT CAV1 complexes. (**A**) Representative 2D class averages of 8S CAV1 oligomers. #, *en face* view; ^, side view; arrow, protruding stalk. Scale bar, 30 nm. (**B**) Cartoon model depicting features and measurements seen in class averages. (**C**) The grayscale intensity around the perimeter of a 2D class average of WT CAV1 as determined using Image J. The positions of the oval used for analysis is shown in white dotted lines on the class average image. The graph shows the grayscale intensity around the complex perimeter. Numbers indicate the position of individual peaks within the profile, presumed to correspond to CAV1 monomers. (**D**) Representative image of negatively stained CAV1-Venus 8S complexes. A schematic of the CAV1-Venus construct is shown above image. Scale bar, 50 nm. (**E**) Representative 2D averages of CAV1-Venus 8S complexes. #, *en face* view; ^, side view; arrow, protruding stalk and Venus tag. Scale bar, 30 nm. (**F**) Representative image of negatively stained MBP-CAV1 8S complexes. A schematic of the MBP-CAV1 construct is shown above image. The dashed white box highlights one 8S MBP-CAV1 disc. Scale bar, 50 nm. (**G**) Examples of individual MBP-CAV1 discs (left) and corresponding cartoon schematics of each particle (right) showing the central disc surrounded by MBP “petals”. Particle with dashed-box is the same complex highlighted in F. Scale bar, 30 nm.

Intriguingly, the outer ring in the *en-face* view of the 2D averages appeared to be composed of distinct globular domains (Figure 2A), although there was variability between classes on how well-defined and how many of these regions were clearly visible. To quantify the size and number of these domains, we measured the intensity of the electron density along the edge of the outer ring of an *en face* 2D average (Figure 2C, Fig S2B). This analysis revealed there are ~10 repeating globular domains that are each ~4.4 nm wide (Figure 2C and S2B, D). We hypothesize these repeating densities may represent CAV1 monomers within the 8S complex.

### The C-terminal region of CAV1 localizes to the stalk region and the N-terminus is localized to the periphery of the discs

There is evidence suggesting the C-terminus of Cav3 is localized to the center of 8S complexes (*18*). However, data supporting this model was principally based on limited labeling by C-terminally directed antibodies (*18*). We therefore sought to directly probe the position of the N- and C-termini of CAV1 within the 8S complexes. For these studies we generated fusions of CAV1 with large tags to serve as fiduciary markers enabling us to trace out the orientation of CAV1 within the complex. We found that addition of C-terminal Venus-tag or a N-terminal MBP-tag (Maltose Binding Protein) was permissive for 8S complex formation (Figure S3A, B) and used these constructs for further studies.

We first examined the C-terminally labeled CAV1-Venus construct. *En-face* views of both individual CAV1-Venus 8S complexes and class averages appeared to be similar to the CAV1 8S complexes in shape (Figure 2A-B and D-E), but had a slighty larger diameter (Figure S2A). Oval plot analysis of the size of the globular domains suggests they contain approximately 8 monomers per complex (Figure S2). Remarkably, an extra density extending from the stalk region was clearly visible in side views of CAV1-Venus complexes (Figure 2D-E). This “fan-shaped” structure emanating from the stalk region of the CAV1-Venus complexes was absent in side-views of CAV1 8S complexes (Figure 2A). These findings suggest the C-terminus of CAV1 is localized to the central stalk region of the 8S complex. Because the C-terminus of CAV1 is known to face the cytoplasm, this additionally suggests that the flat face of the cap is the side of the complex that faces the membrane.

We next evaluated the orientation of the N-terminus of CAV1 using the MBP-CAV1 construct. MBP-CAV1 complexes displayed a sunflower-like pattern, with a central disc structure surrounded by flexible “petals” (Figure 2F-G). The variability of the positions of the MBP densities made it impossible to generate interpretable 2D class averages. Nevertheless, inspection of images of individual MBP-CAV1 8S complexes clearly show that the N-terminus (*i.e.* the MBP) is arrayed around the outer-ring of the 8S disc. The distance and position of the MBP densities relative to the disc varied, suggesting the N-terminus is flexible (Figure 2F-G). Based on measurements of the distances of the MBP-CAV1 complexes seen in Figure 2G, the longest extension of the MBP “petals” in relation to the central discs was ~75 Å. This suggests a possible structural length constraint on the flexible N-terminal region. Finally, the number of flexible “petals” seen around each disc, although difficult to count with precision due to their flexibility, appears to be roughly 8-11 (Figure 2G). This further supports the idea that globular densities that compose the outer-ring of the 8S disc likely each represent one CAV1 monomer (Figure 2C).

### The C-terminus of CAV1 is essential for maintaining the disc shape of the 8S complex

A striking structural feature of the CAV1 8S complex is the protruding C-terminal stalk (Figure 2A-B). To better understand which regions of C-terminus contribute to stalk assembly, we generated a series of C-terminal truncations of CAV1 (Figure 3A). They included constructs designed to disrupt the predicted β-strand located in the extreme C-terminus (*8*). These constructs were expressed in *E. coli*, and the resulting complexes were subsequently purified (Figure S3) and examined by negative stain EM (Figures 3B-C and S4A-B).

**Figure 3.**
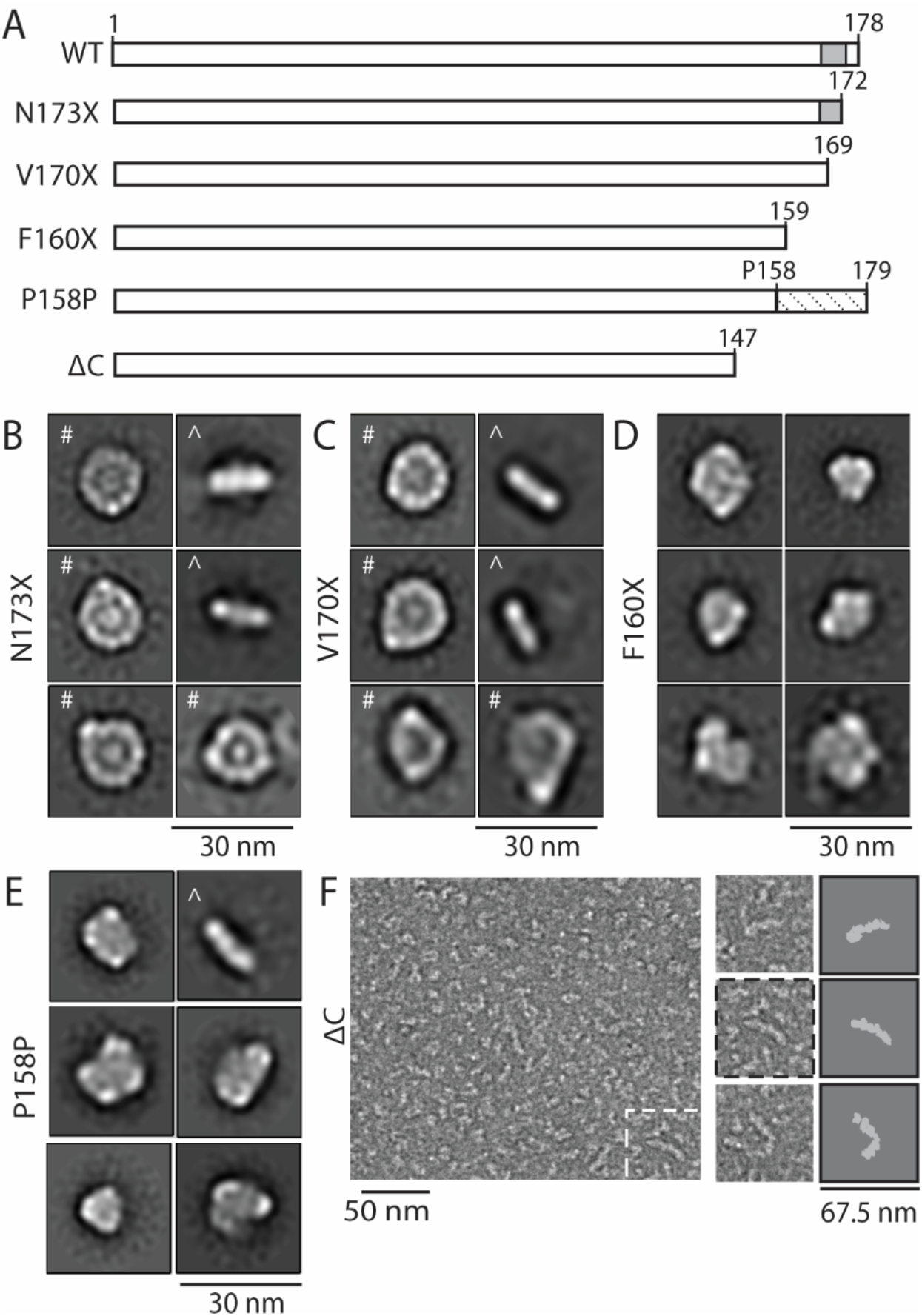
The C-terminus of CAV1 is required for stalk formation and maintaining the disc shape of the 8S complexes. (**A**) Schematic of CAV1 constructs used to test the importance of the C-terminus of CAV1. Grey box, predicted position of β-strand. Dashed box, new C-terminal residues introduced by the P158P frameshift. (**B-E**) Representative class averages of N173X (B), V170X (C), F160X (D), and P158P (E). #, *en face* view; ^, side view. Scale bars, 30 nm. (**F**) Representative negative stain image of CAV1 ΔC (left panel). The dashed white box highlights one linear oligomer. Scale bar, 50 nm. Examples of individual linear ΔC oligomers (middle) and cartoon schematic of each particle (right) are also shown. Particle with dashed-box is the same oligomer highlighted in left panel. Scale bar, 67.5 nm.

The two shortest truncations, N173X and V170X (Figure 3A), formed disc-shaped complexes. *En face* views showed the complexes were generally similar in size and shape as WT CAV1 8S complexes (Figures 3B-C, and S2A). However, N173X contained a smaller protrusion, and the side views of N170X completely lacked a protruding stalk (Figure 3B-C). Furthermore, a fraction of the V170X complexes had a less defined inner ring, and were more heterogenous and less circular than WT CAV1 or N173X complexes (Figure 3C). These results suggest the extreme C-terminal amino acids are required for the generation of the stalk and contribute to the organization of the inner ring region and overall disc-shape of the complex.

To further probe the role of the CAV1 C-terminus in 8S complex organization we examined a naturally occurring disease-associated mutation of CAV1, F160X. Identified in a patient with congenital generalized lipodystrophy (CGL) and pulmonary arterial hypertension (PAH), F160X lacks the last C-terminal 19 amino acids of CAV1 (*22–24*) (Figure 3A). Interestingly, negative stain analysis showed that F160X complexes varied in both size and shape and failed to adopt a uniform disc-like organization or display a clear central stalk (Figures 3D and S4C). Thus, at least a subset of residues 160-178 are required for the creation of the stalk as well as the regularity of the discs.

For comparison, we also examined the structural organization of P158P, another naturally occurring PAH disease mutant (*25, 26*). Unlike F160X which results in a truncation, the frameshift mutation in P158P gives rise to a novel CAV1 C-terminus that is one residue longer than that of WT CAV1 (Figure 3A) (*25, 26*). Interestingly, although the C-termini of P158P and WT CAV1 contain essentially the same number of amino acids (Figure 3A), CAV1-P158P complexes were structurally heterogenous and more similar in appearance to F160X complexes than to WT CAV1 complexes (Figures 2A, 3D-E, and S4C-D). Furthermore, a 2D average of a side view of the P158P complex revealed that the central stalk is missing (Figure 3E, right top panel). This indicates the specific amino acid sequence of residues 158-178 is required to assemble the stalk structure as well as to maintain the correct size and shape of the discs.

We next examined the importance of residues 147-178 in forming CAV1 complexes using a ΔC construct (Figure 3A). The ΔC truncation is predicted to eliminate a portion of a long C-terminal α-helix of CAV1 (residues 132-175) (*27*) as well as a portion of a secondary site identified as important for oligomerization (residues 134-154) (*28*). Negative stain analysis of this construct revealed no disc-like structures; instead we observed heterogenous structures, including some examples of chains of globular domains adopting a “snake-like” appearance (Figure 3F). This suggests residues 1-147 are sufficient to support the assembly of linear oligomers. In addition, since closed discs were seen for F160X but not for ΔC, residues 147-160 appear to contribute to disc closure.

### The N-terminus of CAV1 is also required to generate disc-shaped 8S complexes

We next examined the contributions of the N-terminus to the assembly and structural integrity of 8S complexes. We first studied the naturally occurring β-isoform of CAV1, CAV1β (Figure 4A). CAV1β lacks the first 31 amino acids of the protein, but can assemble into caveolae and is the predominant CAV1 isoform in certain types of mammalian cells (*29*). We examined two CAV1β constructs, one tagged with a 6XHis-tag at the C-terminus and another containing a C-terminal Venus tag. When expressed in *E. coli*, both versions of CAV1β generated 8S-like complexes (Figure S4E-F). The purified CAV1β complexes form discs that contain well-defined inner and outer rings, but are slightly smaller in diameter than observed for full length CAV1 (~13.5 nm versus ~15 nm) (Figures 2A, 4B-C, and S2A; Table S1). The CAV1β-Venus oligomers were more homogeneous than the CAV1β complexes as seen in negative stain 2D averages (Figure 4B-C), suggesting the addition of Venus to the C-terminus may help stabilize the complex. 2D averages of the side view of the complex of CAV1β revealed the presence of a central stalk-like protusion similar to that seen in WT CAV1 complexes, as well as additional fan-shaped density emerging from the stalk for the CAV1β -Venus complexes (Figure 4B-C). Thus, the first N-terminal 31 amino acids of CAV1α, which are missing in CAV1β, are not required for the assembly of the disc-like 8S complex or the central protrusion.

**Figure 4.**
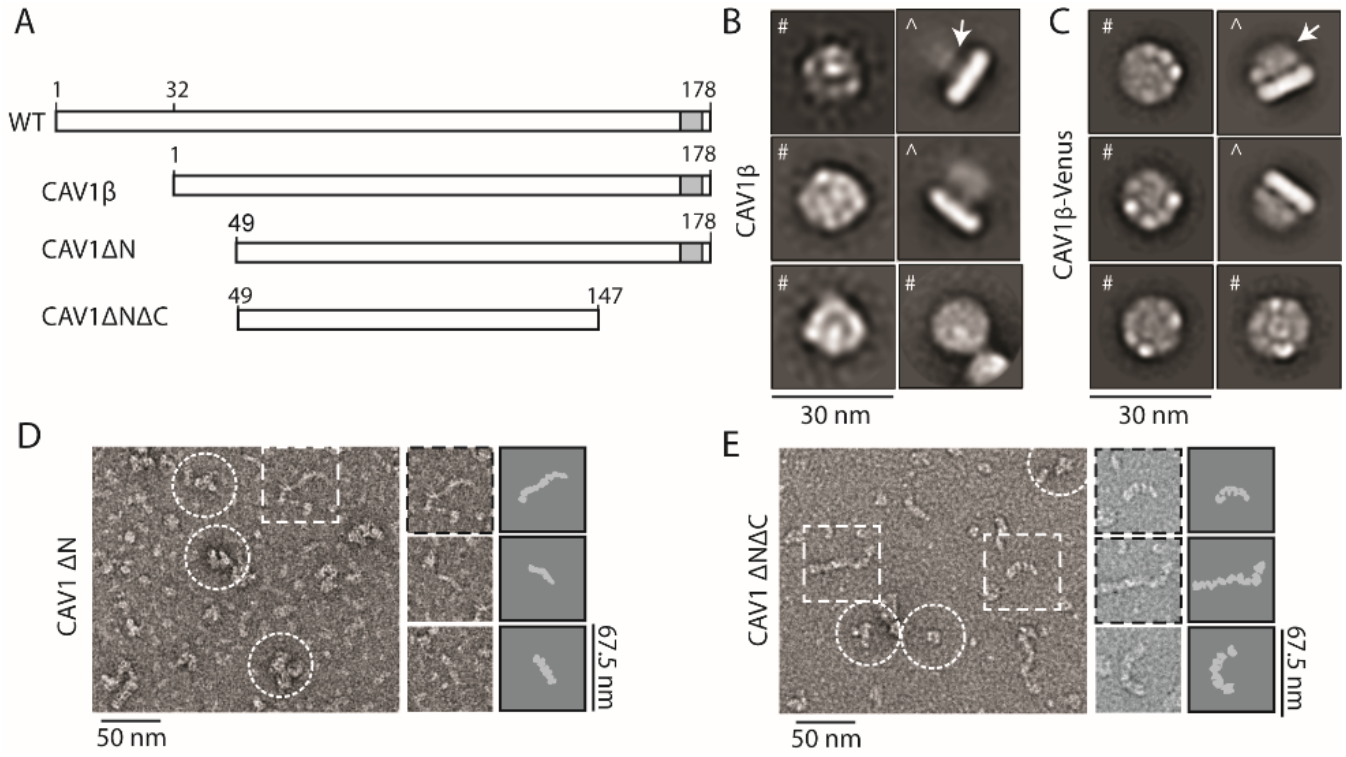
Regions of the N-terminus of CAV1 are required for 8S complex stability. (**A**) Schematic of CAV1 constructs used to test the importance of the N-terminus of CAV1. Grey box, position of predicted β-strand. (B-C) Representative class averages of Cav1β (**B**) and Cav1β-Venus (**C**). #, *en face* view; ^, side view; arrow, protruding stalk. Scale bars, 30 nm. (**D**) Representative negative stain image of CAV1 ΔN (left panel). Dashed white box highlights one linear ΔN oligomer and white circles highlight heterogenous aggregates. Scale bar, 50 nm. Examples of individual linear ΔN oligomers (middle) and corresponding cartoon schematic of each particle (right) are also shown. Particle with dashed-box is the same oligomer highlighted in left panel. Scale bar, 67.5 nm. (**E**) Representative negative stain image of CAV1 ΔNΔC (left panel). Dashed white boxes highlight linear ΔC oligomers and white circles highlight heterogenous aggregates. Scale bar, 50 nm. Examples of individual linear ΔN oligomers (middle) and corresponding cartoon schematic of each particle (right) are also shown. Particles with dashed-box are the same oligomers highlighted in left panel. Scale bar, 67.5 nm.

Residues 1-48 of CAV1 protein are predicted to be structurally disordered (*6*) and are dispensible for caveolae biogenesis to occur in mammalian cells (*4*). To understand how they contribute to the structure of the 8S complex, we next examined a construct lacking the first 48 amino acids of the N-terminal domain, ΔN. Interestingly, ΔN failed to assemble into discs. Instead, it formed a heterogenous mixture of particles, including non-structured protein aggregates of varying size and chains of globular domains of varying lengths with a snake-like appearance (Figure 4D). The snake-like oligomers were generally similar in overall appearance to the structure of the ΔC oligomers (Figures 3F and 4D, right panel). Thus, residues 49-178 can support the assembly of linear and irregular structures, whereas residues 31-48 are additionally required to organize CAV1 oligomers into regular discs.

Given that deletion of either residues 2-48 or 148-178 of CAV1 gave rise to linear oligomers in our 8S complex assembly assay, we wondered whether residues 49-147 are sufficient to support oligomerization. To test this we expressed, purified, and examined a ΔNΔC contruct using negative stain. ΔNΔC retained the ability to generate high molecular weight oligomers as assessed by FPLC (Figure S3). Negative stain analysis showed these correspond to heterogenous linear oligomers that are flexible and vary in size, in addition to smaller oligomeric structures (Figure 4E). We conclude that residues 49-147 are sufficient to support oligomerization, but not for the assembly of fully formed disc-shaped 8S complexes, at least when expressed in *E. coli*.

### 8S complexes are visible in *h*-caveolae

A striking consequence of expressing CAV1 in *E. coli* is the induction of *h*-caveolae (*19*). Previous studies have visualized the distribution of an MBP-tagged form of Cav1 in fixed, detergent extracted *h*-caveolae and revealed a potential polyhedral organization of ring-like Cav1-MBP oligomers (*19*). However, because the structure of isolated CAV1 oligomers themselves was not examined, their positioning within *h*-caveolae could not be directly ascertained (*19*). We speculated that the density of Venus emanating from the C-terminal stalk in the CAV1-Venus complexes, which is predicted to face the cytoplasm, could allow us to visualize the positioning of 8S complexes within *h*-caveolae.

To address this question we purified *h*-caveolae from *E. coli* expressing CAV1-Venus by affinity chromatography and imaged them by negative stain (Figure 5A). The size of *h*-caveolae was heterogenous, with diameters ranging from 17.7 to 54.5 nm and an average diameter of 30.3 ± 8.2 nm (mean ± SD, n = 112) (Figure 5B). Due to this heterogeneity, we were not able to perform averaging across multiple *h*-caveolae. Nevertheless, on the edges of individual *h*-caveolae multiple examples of clustered densities similar to those observed emanating from the central stalk regions of the purified CAV1-Venus complexes could be detected by negative stain EM (Figure 5A and C).

**Figure 5.**
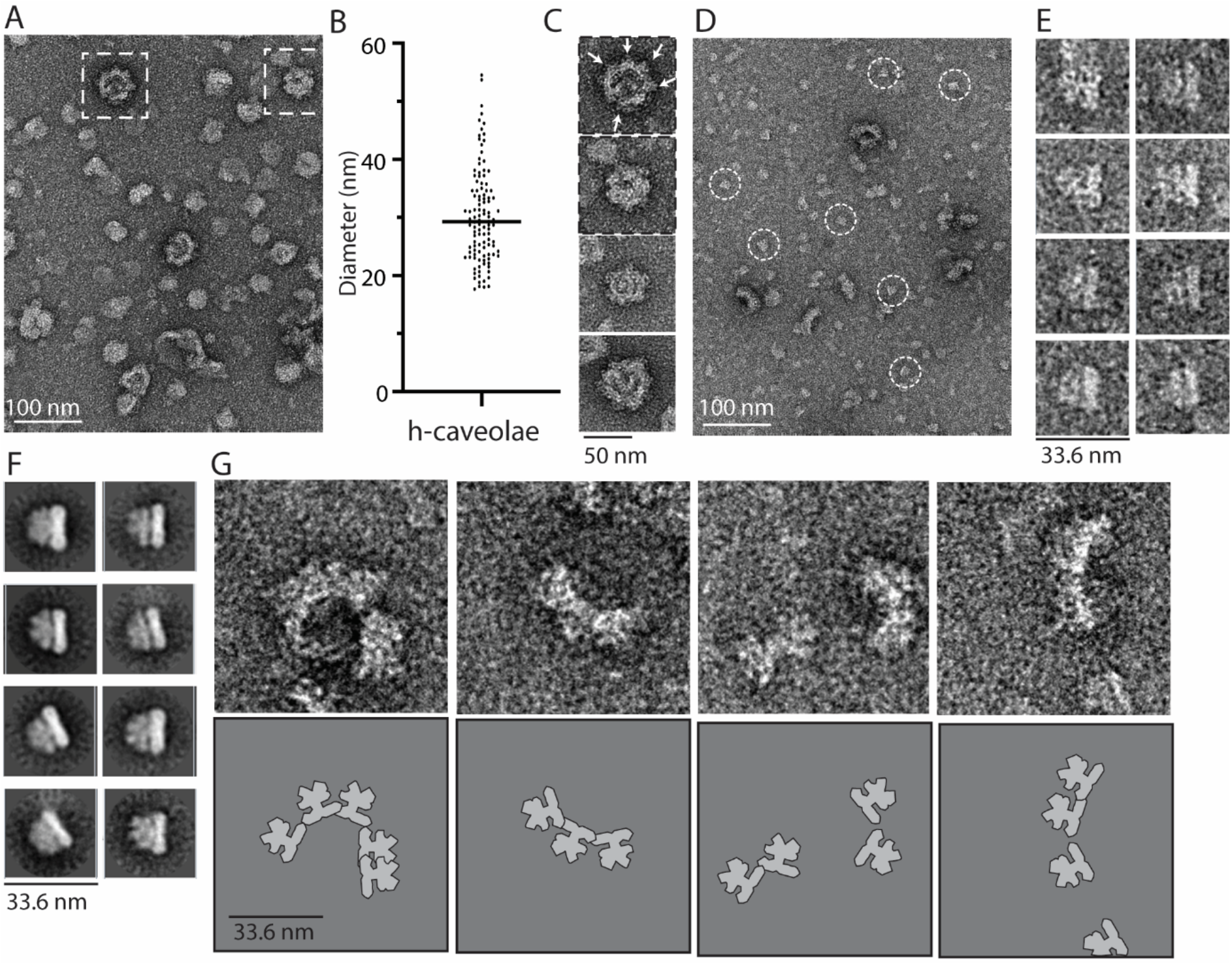
Characterization of *h*-caveolae purified from *E. coli*. (**A**) Representative negative stain image of *h*-caveolae isolated from *E. coli* expressing CAV1-Venus. Scale bar, 100 nm. (**B**) Diameter of 112 individual *h*-caveolae seen in negative stain images. (**C**) Examples of individual *h*-caveolae. Arrows in top panel point to the position of visible 8S CAV1-Venus complexes. Scale bar, 50 nm. (**D**) Representative negative stain image of isolated *h*-caveolae solubilized with 1% Triton-X 100. Examples of individual 8S CAV1-Venus particles are circled in white. Scale bar, 100 nm. (**E**) Examples of individual 8S CAV1-Venus complexes seen in images of solubilized *h*-caveolae. All particles represent side views and have been translationally aligned. Scale bar, 33.6 nm. (**F**) Representative 2D class averages of individual 8S CAV1-Venus complexes obtained from images of Triton-X 100 solubilized *h*-caveolae. All averages represent side views. Scale bar, 33.6 nm. (**G**) Examples of arrays of interconnected 8S Cav1-Venus complexes observed after solubilization with Triton-X 100 (top) and corresponding cartoon representation of the 8S CAV1-Venus complexes in each array (bottom). Scale bar, 33.6 nm.

To determine whether the CAV1-Venus 8S complexes directly interact with one another, we detergent extracted the purified *h*-caveolae using 1% TritonX-100 to remove membrane lipids. This led to the loss of intact *h*-caveolae as expected. Instead we now saw dispersed individual 8S complexes (Figure 5D-E), as well as examples of interconnected 8S complexes that formed arc-like arrays (Figure 5D). 2D class averages of individual 8S complexes obtained from the TritonX-100 extracted sample yielded structures very similar to those obtained for purified 8S-like Cav1-Venus complexes (Figures 2E and 5F). While the arrays were too flexible to generate 2D averages, individual 8S complexes were clearly visible within them (Figure 5G). These results reinforce our findings that these disc-shaped structures correspond to functional complexes that represent the fundamental building blocks of *h*-caveolae.

## Discussion

### Overall organization and novel structural features of the 8S CAV1 complex

Despite their importance in caveolae biogenesis, the structural organization of 8S complexes formed by the monotopic membrane protein caveolin has remained enigmatic. Here we provide critical new insights into the architecture and assembly of recombinant human CAV1 8S complexes using single particle electron microscopy.

We show that detergent-solubilized, negatively-stained 8S CAV1 complexes isolated from *E. coli* are disc shaped and exhibit a toroidal organization consisting of an outer ring of globular densities surrounding a weaker density region, and a higher density central structure (Figure 2A-B). The overall shape of the complex is thus consistent with previous studies suggesting CAV1 has a intrinsic propensity to form disc- or ring-like structures (*2, 13*). Strikingly, many of the features of the CAV1 8S complex are very similar to those previously reported for negatively stained Cav3 complexes isolated from insect cells (*18*), including their overall dimensions (15 nm × 4.8 nm for CAV1 versus 16.5 nm × 5.5 nm for Cav3). A centrally located protruding structure is also apparent in both complexes. For the case of CAV1, this corresponds to a stalk-like protrusion in side views (Figure 2A), whereas for Cav3 the central structure appears to be more cone-shaped (*18*). The fact that so many similarities exist between these structures, despite having been isolated for two different caveolin family members from two different expression systems, strongly suggests this represents the characteristic architecture of 8S complexes.

We hypothesize the repeating globular units comprising the outer ring of the 8S complexes represent Cav1 monomers. In our experiments, the number of monomers per complex ranged from 8 to 11 by counting the number of discrete densities and estimates of monomer size relative to that of the 8S complex itself. For example, assuming the average diameter for WT CAV1 8S complexes is 14.4 nm and the average distance each monomer occupies along the circumference of the complex is 4.4 nm (Figure S2), we estimate each WT CAV1 complex contains ~10 monomers. Using a similar approach, each CAV1-Venus 8S complex would be predicted to contain ~8 monomers. These values are within range of those previous estimates from biochemical studies as well as the report that negatively stained Cav3 complexes contain 9 monomers (*18*). Interestingly, even for complexes containing WT CAV1, packing of globular domains of the protein in the outer ring of the 8S complexes was not regular across the entire ring; in several cases, globular domains appeared to protrude from the ring, breaking the symmetry of the structure. It is thus possible that structural heterogeneity is a native feature of the complexes.

Our results also enable us to pinpoint the position of the N- and C-termini within the complex. We show the N-terminus of CAV1 localizes on the periphery of the outer ring and confirm it assumes a flexible conformation based on the irregular distribution of the MBP tag (Figure 2F-G). This is consistent with reports that the N-terminal region of CAV1 is unstructured (*6*) and the first 48 residues in particular are protease sensitive (and thus solvent accessible) in *h*-caveolae (*19*). This also strongly suggests that interactions between adjacent discs to generate higher-order oligomers, such as are expected to occur in caveolae, are mediated by the N-terminal domain of CAV1. We also provide definitive evidence that the C-terminal domain of CAV1 localizes to the center of the complex and show it contributes to the assembly of the central stalk (Figures 2D-E and 3B-C). The resolution of the negative stain structure is too low to definitively determine the secondary structure of this region of the protein, but we speculate this could be a β-barrel given the predicted propensity of the extreme C-terminal domain to form a beta strand (*8*). Our findings also suggest interactions can occur between stalks, as evidenced by the presence of dimers of WT CAV1 (Figure 1). It remains to be determined whether this is a physiologically relevant interaction. Clearly, obtaining higher resolution structural information, such as from cryoEM, will be essential to pinpoint the exact positioning of these and other other key structural elements of CAV1 within the complex. It will also be important to examine the structure of 8S complexes in membrane environments, including in the presence of cholesterol, a key regulator of caveolae structure.

Based on the positions of the C- and N-terminus of CAV1 within the 8S complex, we can infer where other domains of the protein likely reside. The flat face of the complex, opposite of that from which the C-terminal stalk emerges, can be assigned as the membrane associated surface of the complex given that the C-terminus of CAV1 is known to face the cytoplasm. Since the N-terminus is localized at the periphery of the disc, it seems likely other domains found in the N-terminal half of the protein, including the oligomeriation and scaffolding domains, also reside in the outer ring. The central location of the C-terminus implies the three palmitoylation sites of CAV1 (Cys133, Cys143, and Cys156) are likely positioned in the center of the complex. This would generate a very high density of palmitates in this area, which in turn could potentially generate a highly ordered nanodomain.

### Proposed model for how 8S complexes assemble and how defects in assembly contribute to disease

To better understand the structural basis for the assembly of 8S complexes, we probed the contributions of the N- and C-terminal domains of CAV1 to their overall organization (Figures 3-4). The naturally occuring CAV1β isoform, which lacks residues 1-31, generated disc shaped structures similar in shape but slightly smaller than those formed by WT CAV1α (Figures 4B-C and S2A). In contrast, truncation of residues 1-47 of the N-terminus more severely disrupted the size and shape of complexes, leading to the assembly of irregular aggregates and linearized chains of globular domains (Figure 4D). Truncations of the extreme C-terminus (residues 170-178) maintained overall disc structure but caused a loss of the central protrusion and some rearrangements within the central region of the complex (Figure 3B-C), whereas disease-associated mutants of the protein (F160X and P158P) generated irregular discs or aggregates (Figure 3D-E). Further truncation of the C terminus, either in the presence (ΔC) or absence (ΔNΔC) of an intact N-terminus, ultimately led to the formation of chains of globular domains rather than toroidal discs (Figures 3F and 4E).

Taken together, these findings suggest a potential mechanism for how assembly of disc-shaped 8S complexes occurs (Figure 6). In this working model, CAV1 monomers can be thought of as wedge-shaped “sections” that come together to form an “orange slice” (Figure 6A). One important set of interactions between the “wedges” occurs at the outer ring of the disc, as revealed when the C-terminal domain of CAV1 is truncated (Figure 6B). We speculate that this oligomerization event is driven by the core region of the protein (residues 49-147). To support such a packing arrangement, heterotypic interactions involving at least two surfaces of each monomer would be required (Figure 6C). While the identity of these interaction surfaces is currently unknown, the well-studied oligomerization domain likely serves as one interaction interface. Additional binding events between monomeric “wedges” occur within the inner ring of the complex where the C-terminal domain of CAV1 is located (Figure 6C). This includes interactions between the extreme C-terminal domains of CAV1 to create the central tubular stalk structure, possibly via the formation of a β-barrel as discussed above. We speculate the formation of this tubular structure may not only serve as a mechanism that helps stabilize the the discs but also could help define the ultimate size the discs can attain. The N-terminal domain of CAV1 also aids in the generation of regular discs. Residues 32-48 appear to be especially important in this process, since deletion of residues 1-31 support native-like complex assembly whereas deletion of residues 2-48 do not. While the structural basis for this activity is currently unknown, it is interesting to note that residues 30-50 have previously been predicted to generate an amphipathic alpha helix (*30*).

**Figure 6.**
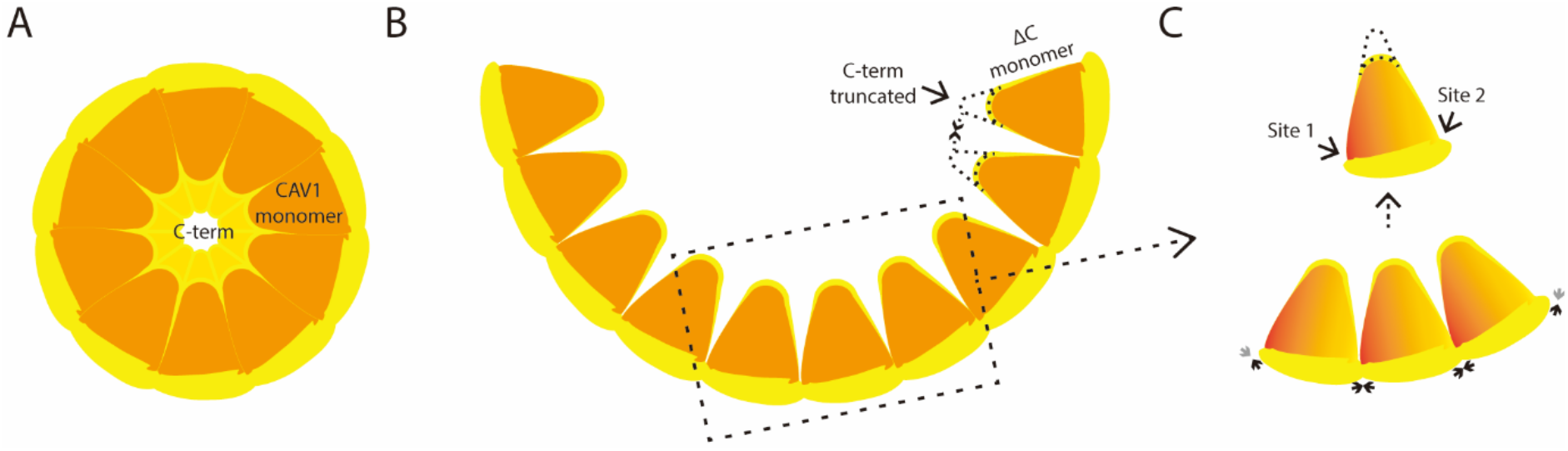
“Orange” model of 8S complex assembly. (**A**) Cav1 monomers form wedge-like “sections” that self assemble into a disc-shaped complex containing inner and outer rings depicted here as an orange slice. In this model, the C-terminus of Cav1 is localized to the center of the complex whereas the N-terminus is found on the periphery of the disc. Interactions between monomers that contribute to disc formation occur both in the outer ring and inner ring. (**B**) In the absence of an intact C-terminus, Cav1 assembles into linear oligomers stabilized by interactions between adjacent monomers in the outer ring. (**C**) Along the outer ring, interactions between Cav1 monomers are predicted to occur at two binding interfaces, as opposed to primarily involving homotypic interactions between oligomerization domains of the monomers.

The “orange” model of 8S complex assembly has several interesting implications. First, the organization of wedge-shaped monomers into a disc gives rise to a structure that could potentially incorporate varying numbers of n-mers. Second, this model suggests it is unlikely that homotypic interactions between the oligomerization domains of adjacent Cav1 monomers occurs, as was suggested in earlier models (*31*). Instead, it predicts the oligomerization domain undergoes heterotypic interactions with a secondary binding site of the protein. Third, the centrally located C-terminus contributes importantly to the formation of the stalk and inner ring in this model. This can help explain why progressive truncations of the C-terminus increasingly destabilize Cav1 complexes (*15*). Finally, the model predicts that monomers must ultimately assemble into a closed ring to form a mature 8S complex. Although the exact mechanism by which this structure is generated remains to be determined, it is not difficult to imagine that it occurs via a cooperative process (*15*).

Our results also shed light on how the human CAV1 mutations F160X and P158P contribute to disease. We previously reported both mutants are capable of generating 8S complexes when expressed either individually or in combination with WT CAV1 (*22, 25*). Interestingly, the complexes containing F160X are destabilized compared to those formed by WT CAV1 (*22*). Our current findings suggest this destabilization is the direct result of disruption of the central stalk together with packing defects of the protein in the center of the 8S complex resulting from the loss of residues 160-178 due to the frameshift. This model further predicts the C-terminus of WT CAV1 would also fail to become incorporated into the protrusion when co-expressed with the mutant proteins. Consistent with this, we found the C-terminus of CAV1 is aberrantly exposed in caveolae in patient fibroblasts heterozygous for the F160X mutant as well as when co-expressed with WT CAV1 in Cav1^−/−^ mouse embryonic fibroblasts (*22*). Patient cells expressing a copy of either F160X or P158P together with a copy of WT CAV1 also exhibit other abnormalities, including decreased co-localization of cavin with CAV1 in presumptive caveolae and decreased DRM association of CAV1 (*22, 25*). Our current findings implicate defects in the central region of the 8S complexes as giving rise to these irregularities.

### Implications for how 8S complexes pack in caveolae and *h*-caveolae

Current models suggest the polygonal shape of mammalian caveolae is generated by a coat complex consisting of an inner polyhedral cage formed by Cav1 surrounded by an outer filamentous layer composed of cavins (*2, 32*). The presence of the cytoplasmic coat of cavins has made it difficult to gain further insights into the organization of 8S complexes within this structure. In contrast, the absence of cavins in *h*-caveolae offers the potential to directly visualize Cav1 embedded in membrane under conditions where it induces membrane curvature. Here, we were able to directly observe multiple 8S complexes in purified negatively stained *h*-caveolae with the help of Venus as a fiduciary marker (Figure 5A-C). We also detected individual 8S-like complexes in preparations of purified, detergent-solubilized *h*-caveolae (Figure 5D-F). Examples of groups of two or three conjoined 8S complexes aligned in a side-to-side organization were also apparent in preparations of detergent solubilized, purified *h*-caveolae (Figure 5D and G). Thus, *h*-caveolae do in fact appear to be composed of multiple copies of these discs. The monomeric, dimeric, and trimeric forms of the 8S complexes we observed could potentially correspond to the 200 kDa, 400 kDa, and 600 kDa species detected in original descriptions of caveolin oligomers (*12*). More work will be required to determine exactly how Cav1 bends membranes to form *h*-caveolae and caveolae, and whether the same or different mechanisms are responsible for mammalian caveolae biogenesis and *h*-caveolae assembly. In this regard, it is interesting to note that ectopic overexpression of other membrane proteins can also generate intracellular membranes in *E. coli,* indicating multiple mechanisms may be at play in this process (*33*).

### Caveolin as a model for understanding monotopic membrane protein structure

In addition to serving as a key building block of caveolae, caveolin is a member of the poorly understood family of monotopic membrane proteins. Unlike traditional transmembrane proteins, monotopic membrane proteins contain structural elements that enter and exit on a single face of the membrane as opposed to completely spanning the membrane bilayer. These proteins have been particularly recalcitrant for structural studies, as there are only a few with available structures (*10, 34, 35*). The vast majority of the monotopic membrane proteins with known structures are enzymes that share homology with soluble proteins that catalyze similar reactions. For these proteins, membrane attachment facilitates access to their membrane-bound substrates. These monotopic enzymes are typically assembled as either monomers or dimers (*10*). One exception is seipin, a dodecameric structural protein involved in biogenesis of lipid droplets (*10, 35*). The caveolin family members Cav1 and Cav3 thus constitute additional examples of structural monotopic proteins that function as higher-ordered oligomers. Most of the monotropic membrane proteins shallowly associate with membranes through amphipathic helices positioned parallel to the membrane plane, hydrophobic loops extending into the membrane core, and/or hydrophobic patches along with hydrophilic residues that interact with phospholipid head groups. Alternatively, some penetrate more deeply into the membrane through reentrant helical domains, disrupting the lipid bilayer (*36*). Intriguingly, Cav1 is thought to not only contain a U-shaped helix-break-helix intramembrane domain, but is also predicted to contain one or more amphipathic helices (*6*). Thus, further structural analysis of Cav1 holds the potential to offer critical insights into the mechanisms by which monotopic proteins associate with biological membranes and assemble into higher order complexes to fulfill unique structural roles.

## Materials and Methods

### Cloning

A summary of the constructs used in this study is provided in Table S2, and the primers used to generate them are described in Table S3. Histidine tagged truncations and disease-associated mutants of CAV1 were generated by PCR from a pET24-CAV1-6His plasmid (a gift from Dr. K. Jebrell Glover, Lehigh University) containing a cDNA sequence of WT human CAV1 codon optimized for *E.coli* expression and cloned into a pET20 vector (Novagen No. 69739) using Nde I and Xho I restriction sites. To generate MBP-CAV1, the cDNA of human CAV1 was PCR amplified from the mammalian expression plasmid Emerald-CAV1 C-10 (Addgene No. 54025, a gift from Dr Michael Davidson) and cloned into a pNMTMA-MBP-Cav1 (canine) plasmid described in (*19*) (a gift from Dr. Robert Parton, University of Queensland) using BamH I and EcoR I restriction sites to replace the sequence of canine Cav1. To better control the expression, the human CAV1 fragment was further moved into a pET28-MBP-TEV plasmid (Addgene No. 69929) described in (*37*) using BamH I and Xho I restriction sites. CAV1-Venus was synthesized and subcloned into the above mentioned pET28 vector by GenScript (Piscataway, NJ). CAV1β-Venus was generated from the CAV1-Venus construct by introducing flanking NcoI and HindIII sites. After the restriction digest, the insert was ligated into the mutated pET28/Tev-mVenus-His vector, carrying a HindIII site which was added by QuikChange Lightning Site-Directed Mutagenesis (Agilent). Following the successful ligation, the HindIII site was destroyed to restore the original sequence. All constructs were confirmed by sequencing.

### Expression of CAV1 and purification of CAV1 complexes and *h*-caveolae

Caveolin proteins were expressed in E.*coli* BL21 using the auto-induction expression system (*38*). Specifically, monoclonal bacteria were first cultured in non-inducing MDG media at 37°C, 250 rpm for 20 h, then the cultures were enlarged in auto-inducing ZYM-5052 media at 25°C, 350 rpm for 24 h. E.*coli* cells were resuspended with buffer (200 mM NaCl, 20 mM Tris-HCl pH 8.0) and homogenized with pressure homogenizer (EmulsiFlex^®^-C3 or French Press); 1 mM DTT and PMSF were added just before homogenization. Large cell debris was removed by 15 min centrifugation at 9,000 rpm and 4°C. Total membranes were pelleted at 40,000 rpm (Ti-50.2 rotor) at 4°C for 1 h, and then were homogenized in a buffer composed of 200 mM NaCl, 20 mM Tris-HCl pH 8.0 and 1mM DTT using a Dounce tissue grinder. Solubilization of 8S complexes was performed by mixing the homogenized sample at 4°C for 2 hours after addition of 2% C12M (Anatrace). After removing the insoluble material by centrifugation at 42,000 rpm (Ti-45 rotor) for 35 min, CAV1 was purified using nickel sepharose-based affinity purification. Elutions with CAV1 were concentrated and further purified by size exclusion chromatography using Superpose 6 Increase 10/300 GL (GE Healthcare, Illinois) equilibrated with a buffer composed of 200 mM NaCl, 20 mM Tris-HCl pH 8.0, 1mM DTT and 0.05% C12M.

For *h-*caveolae purification, the homogenized total membranes were directly subjected to the nickel sepharose-based purification procedure without prior detergent extraction. Where indicated, the purified *h*-caveolae were subsequently solubilized with 1% TX-100 on a rotator at 4°C for 30 min.

### Electrophoresis and western blotting

Electrophoresis (SDS–PAGE and Blue native) and Western blotting were performed as described previously (*20, 22, 25*). Rabbit anti-Cav1 polyclonal antibody (catalog number 610059) from BD Biosciences (San Jose, CA) was used with 1:10,000 times dilution.

### Negative stain preparation and electron microscopy

Samples of purified Cav1 constructs were prepared for negative stain EM using established methods (*39*). In brief, 200-mesh copper grids covered with carbon-coated collodion film (EMS, Hatfield, PA, USA) were glow discharged for 30 s at 10 mA in a PELCO easiGlow™ glow discharge unit (Fresno, CA, USA). Aliquots (3.5 μl) of the Cav1 purifications were adsorbed to the grids and incubated for 1 minute at room temperature. Samples were then washed with 2 drops of water and stained with 2 successive drops of 0.7% (w/v) uranyl formate (EMS, Hatfield, PA, USA) followed by blotting until dry. Samples were visualized on a Tecnai Spirit T12 transmission electron microscope equipped with a field emission gun operating at an accelerating voltage of 120 keV (Thermo Fisher Scientific Inc., Waltham, MA, USA) at a nominal magnification of 26,000x (2.34 Å per pixel) or using a Thermo Fisher Tecnai T20 equipped with a field emission gun operating at an accelerating voltage of 200 keV. For the CAV1-Venus sample low-dose data was collected using Leginon software (*21*) on a 4k × 4k CCD Ultrascan camera (Gatan, Pleasanton, CA) at −1.5 μm defocus value. For all the other CAV1 samples data was collecting using Leginon software on a 4k × 4x Rio CMOS camera (Gatan, Pleasanton, CA) at −1.5 μm defocus value.

Purified intact *h*-caveolae and detergent-solublized *h*-caveolae were prepared for negative-stain electron microscopy by standard methods. An aliquot was applied to glow-discharged continuous carbon grids for 1 minute, blotted, washed with 3 drops water then 3 drops of 2% uranyl acetate, with the final drop incubating for 1 minute before blotting to dryness. The grids were imaged in an F20 electron microscopy (Thermo-Fisher, formerly FEI) operating at 120kV at magnifications ranging from 29,000X-62,000X, with images at 50,000x and a pixel size of 2.25Å used for image processing.

### Image processing and particle classification

Negative stain images were manually curated and images were processed using Relion 3.0.8 (*21*). For each dataset approximately 1,000 particles were manually picked and subjected to 2D classification. Several of the highest resolution classes were then used as references for particle selection on all images. Particles were then extracted into 128 pixel boxes (30 nm × 30 nm) followed by 2D classification. Datasets include 16,834 particles for WT CAV1, 11,016 particles for CAV1-Venus, 15,106 for CAV1 V170X, 13,012 particles for CAV1 N173X, 37,568 particles for CAV1β-Venus and 18,424 particles for CAV1β.

### Data analysis and diameter measurements

Oval plot analysis was conducted according to the method shared by dscho on ImageJ forum: http://imagej.1557.x6.nabble.com/Circular-plot-with-ImageJ-td3687397.html. All diameter measurements were carried out using the circular “*en-face*” views only. Images were downloaded from the Leginon web interface (*21*) in JPG format and opened in ImageJ (*40*). To perform diameter measurements, a spatial scale of the active image was defined by measuring a 200 nm scale bar on a single micrograph with the *set scale* function. Diameter measurements for each dataset were made using a line tool followed by the *measure* function. To obtain statistically robust results, 200 measurements were made for each dataset. Prism 8 (v8.4.2; GraphPad Software, San Diego, CA, USA) was used to plot the results and perform statistical analyses.

## Abbreviations

Cav: Caveolin
EM: electron microscopy
*h*-caveolae: heterologous caveolae
WT: wild type
MBP: Maltose binding protein
CGL: congenital generalized lipodystrophy
PAH: pulmonary arterial hypertension
FPLC: Fast protein liquid chromatography

## Acknowledgements

We thank Dr. Robert Parton and Dr. K. Jebrell Glover for providing plasmids. We also thank Margaret Elmer-Dixon and members of the Ohi labs for helpful discussions and Ting Wang for editorial assistance with an early draft of the manuscript. The University of Michigan Cryo-EM Facility (U-M Cryo-EM) has received generous support from the U-M Life Sciences Institute and the U-M Biosciences Initiative. The University of Virginia Molecular Electron Microscopy Core facility at the University of Virginia is supported in part by the School of Medicine and built with NIH grant G20-RR31199. We acknowledge M. Su, A. Bondy, L. Keopping, C. Lilienthal, B. Battey, and L. Chang for technical support. This work was supported by NIH grants R01HL144131 (M.D.O and A.K.K.) and S10OD020011 (LSI computation).

## Author Information

### Contributions

E.K., D.P.C., H.S.M., and B.H. developed caveolin-1 purification procedures. B.H. and Y.P. cloned and biochemically purified all Cav1 constructs. J.C.P., J.L.H., E.B., and K.D. collected and processed the EM data. M.D.O. and A.K.K. conceived the project and provided feedback on all experiments and data processing. B.H., J.C.P., J.L.H, M.D.O, and A.K.K. wrote the initial drafts of the manuscript. All authors interpreted the data, discussed the results, and provided feedback on the final version of the manuscript.

## References

1. R. G. Parton, M. A. del Pozo, Caveolae as plasma membrane sensors, protectors and organizers. Nat Rev Mol Cell Biol 14, 98–112 (2013).

2. M. Stoeber, P. Schellenberger, C. A. Siebert, C. Leyrat, A. Helenius, K. Grunewald, Model for the architecture of caveolae based on a flexible, net-like assembly of Cavin1 and Caveolin discs. Proc Natl Acad Sci U S A 113, E8069–E8078 (2016).

3. C. Lamaze, N. Tardif, M. Dewulf, S. Vassilopoulos, C. M. Blouin, The caveolae dress code: structure and signaling. Curr Opin Cell Biol 47, 117–125 (2017).

4. M. Kirkham, S. J. Nixon, M. T. Howes, L. Abi-Rached, D. E. Wakeham, M. Hanzal-Bayer, C. Ferguson, M. M. Hill, M. Fernandez-Rojo, D. A. Brown, J. F. Hancock, F. M. Brodsky, R. G. Parton, Evolutionary analysis and molecular dissection of caveola biogenesis. J Cell Sci 121, 2075–2086 (2008).

5. B. Sinha, D. Koster, R. Ruez, P. Gonnord, M. Bastiani, D. Abankwa, R. V. Stan, G. Butler-Browne, B. Vedie, L. Johannes, N. Morone, R. G. Parton, G. Raposo, P. Sens, C. Lamaze, P. Nassoy, Cells respond to mechanical stress by rapid disassembly of caveolae. Cell 144, 402–413 (2011).

6. K. T. Root, J. A. Julien, K. J. Glover, Secondary structure of caveolins: a mini review. Biochem Soc Trans 47, 1489–1498 (2019).

7. K. T. Root, S. M. Plucinsky, K. J. Glover, Recent progress in the topology, structure, and oligomerization of caveolin: a building block of caveolae. Curr Top Membr 75, 305–336 (2015).

8. N. Ariotti, J. Rae, N. Leneva, C. Ferguson, D. Loo, S. Okano, M. M. Hill, P. Walser, B. M. Collins, R. G. Parton, Molecular characterization of caveolin-induced membrane curvature. J Biol Chem 290, 24875–24890 (2015).

9. J. Lee, K. J. Glover, The transmembrane domain of caveolin-1 exhibits a helix-break-helix structure. Biochim Biophys Acta 1818, 1158–1164 (2012).

10. K. N. Allen, S. Entova, L. C. Ray, B. Imperiali, Monotopic Membrane Proteins Join the Fold. Trends Biochem Sci 44, 7–20 (2019).

11. M. Sargiacomo, P. E. Scherer, Z. Tang, E. Kubler, K. S. Song, M. C. Sanders, M. P. Lisanti, Oligomeric structure of caveolin: implications for caveolae membrane organization. Proc Natl Acad Sci U S A 92, 9407–9411 (1995).

12. S. Monier, R. G. Parton, F. Vogel, J. Behlke, A. Henske, T. V. Kurzchalia, VIP21-caveolin, a membrane protein constituent of the caveolar coat, oligomerizes in vivo and in vitro. Mol. Biol. Cell 6, 911–927 (1995).

13. I. Fernandez, Y. Ying, J. Albanesi, R. G. Anderson, Mechanism of caveolin filament assembly. Proc Natl Acad Sci U S A 99, 11193–11198 (2002).

14. A. Hayer, M. Stoeber, C. Bissig, A. Helenius, Biogenesis of caveolae: stepwise assembly of large caveolin and cavin complexes. Traffic 11, 361–382 (2010).

15. X. Ren, A. G. Ostermeyer, L. T. Ramcharan, Y. Zeng, D. M. Lublin, D. A. Brown, Conformational defects slow Golgi exit, block oligomerization, and reduce raft affinity of caveolin-1 mutant proteins. Mol Biol Cell 15, 4556–4567 (2004).

16. A. R. Busija, H. H. Patel, P. A. Insel, Caveolins and cavins in the trafficking, maturation, and degradation of caveolae: implications for cell physiology. Am J Physiol Cell Physiol 312, C459–C477 (2017).

17. Z. Tang, P. E. Scherer, T. Okamoto, K. Song, C. Chu, D. S. Kohtz, I. Nishimoto, H. F. Lodish, M. P. Lisanti, Molecular cloning of caveolin-3, a novel member of the caveolin gene family expressed predominantly in muscle. J Biol Chem 271, 2255–2261 (1996).

18. G. Whiteley, R. F. Collins, A. Kitmitto, Characterization of the molecular architecture of human caveolin-3 and interaction with the skeletal muscle ryanodine receptor. J Biol Chem 287, 40302–40316 (2012).

19. P. J. Walser, N. Ariotti, M. Howes, C. Ferguson, R. Webb, D. Schwudke, N. Leneva, K. J. Cho, L. Cooper, J. Rae, M. Floetenmeyer, V. M. Oorschot, U. Skoglund, K. Simons, J. F. Hancock, R. G. Parton, Constitutive formation of caveolae in a bacterium. Cell 150, 752–763 (2012).

20. B. Han, A. Tiwari, A. K. Kenworthy, Tagging strategies strongly affect the fate of overexpressed caveolin-1. Traffic 16, 417–438 (2015).

21. C. Suloway, J. Pulokas, D. Fellmann, A. Cheng, F. Guerra, J. Quispe, S. Stagg, C. S. Potter, B. Carragher, Automated molecular microscopy: the new Leginon system. J Struct Biol 151, 41–60 (2005).

22. B. Han, C. A. Copeland, Y. Kawano, E. B. Rosenzweig, E. D. Austin, L. Shahmirzadi, S. Tang, K. Raghunathan, W. K. Chung, A. K. Kenworthy, Characterization of a caveolin-1 mutation associated with both pulmonary arterial hypertension and congenital generalized lipodystrophy. Traffic 17, 1297–1312 (2016).

23. A. Garg, M. Kircher, M. Del Campo, R. S. Amato, A. K. Agarwal, Whole exome sequencing identifies de novo heterozygous CAV1 mutations associated with a novel neonatal onset lipodystrophy syndrome. Am J Med Genet A 167, 1796–1806 (2015).

24. I. Schrauwen, S. Szelinger, A. L. Siniard, A. Kurdoglu, J. J. Corneveaux, I. Malenica, R. Richholt, G. Van Camp, M. De Both, S. Swaminathan, M. Turk, K. Ramsey, D. W. Craig, V. Narayanan, M. J. Huentelman, A frame-shift mutation in CAV1 is associated with a severe neonatal progeroid and lipodystrophy syndrome. PLoS One 10, e0131797 (2015).

25. C. A. Copeland, B. Han, A. Tiwari, E. D. Austin, J. E. Loyd, J. D. West, A. K. Kenworthy, A disease-associated frameshift mutation in caveolin-1 disrupts caveolae formation and function through introduction of a de novo ER retention signal. Mol Biol Cell 28, 3095–3111 (2017).

26. E. D. Austin, L. Ma, C. LeDuc, E. Berman Rosenzweig, A. Borczuk, J. A. Phillips, 3rd, T. Palomero, P. Sumazin, H. R. Kim, M. H. Talati, J. West, J. E. Loyd, W. K. Chung, Whole exome sequencing to identify a novel gene (caveolin-1) associated with human pulmonary arterial hypertension. Circ Cardiovasc Genet 5, 336–343 (2012).

27. S. M. Plucinsky, K. J. Glover, Secondary structure analysis of a functional construct of caveolin-1 reveals a long C-terminal helix. Biophys J 109, 1686–1688 (2015).

28. T. Machleidt, W. P. Li, P. Liu, R. G. Anderson, Multiple domains in caveolin-1 control its intracellular traffic. J. Cell Biol. 148, 17–28 (2000).

29. T. Fujimoto, H. Kogo, R. Nomura, T. Une, Isoforms of caveolin-1 and caveolar structure. J Cell Sci 113 Pt 19, 3509–3517 (2000).

30. R. G. Parton, M. Hanzal-Bayer, J. F. Hancock, Biogenesis of caveolae: a structural model for caveolin-induced domain formation. J Cell Sci 119, 787–796 (2006).

31. K. S. Song, Z. L. Tang, S. W. Li, M. P. Lisanti, Mutational analysis of the properties of caveolin-1. A novel role for the C-terminal domain in mediating homo-typic caveolin-caveolin interactions. J. Biol. Chem. 272, 4398–4403 (1997).

32. A. Ludwig, B. J. Nichols, S. Sandin, Architecture of the caveolar coat complex. J Cell Sci 129, 3077–3083 (2016).

33. N. Jamin, M. Garrigos, C. Jaxel, A. Frelet-Barrand, S. Orlowski, Ectopic neo-formed intracellular membranes in Escherichia coli: a response to membrane protein-induced stress involving membrane curvature and domains. Biomolecules 8, (2018).

34. L. C. Ray, D. Das, S. Entova, V. Lukose, A. J. Lynch, B. Imperiali, K. N. Allen, Membrane association of monotopic phosphoglycosyl transferase underpins function. Nat Chem Biol 14, 538–541 (2018).

35. X. Sui, H. Arlt, K. P. Brock, Z. W. Lai, F. DiMaio, D. S. Marks, M. Liao, R. V. Farese, Jr., T. C. Walther, Cryo-electron microscopy structure of the lipid droplet-formation protein seipin. J Cell Biol 217, 4080–4091 (2018).

36. K. Balali-Mood, P. J. Bond, M. S. Sansom, Interaction of monotopic membrane enzymes with a lipid bilayer: a coarse-grained MD simulation study. Biochemistry 48, 2135–2145 (2009).

37. H. Currinn, B. Guscott, Z. Balklava, A. Rothnie, T. Wassmer, APP controls the formation of PI(3,5)P(2) vesicles through its binding of the PIKfyve complex. Cell Mol Life Sci 73, 393–408 (2016).

38. F. W. Studier, Protein production by auto-induction in high density shaking cultures. Protein Expr Purif 41, 207–234 (2005).

39. M. Ohi, Y. Li, Y. Cheng, T. Walz, Negative staining and image classification - powerful tools in modern electron microscopy. Biol Proced Online 6, 23–34 (2004).

40. C. A. Schneider, W. S. Rasband, K. W. Eliceiri, NIH Image to ImageJ: 25 years of image analysis. Nat Methods 9, 671–675 (2012).

