## Supplementary Material for "Structure and assembly of CAV1 8S complexes revealed by single particle electron microscopy"

**Table S1. Summary of 8S complex diameter measurements (accompanies Fig. S2A)**

| | WT CAV1 | CAV1-Venus | N173X | V170X | CAV1β-Venus | CAV1β |
|---|---|---|---|---|---|---|
| N | 200 | 200 | 200 | 200 | 200 | 200 |
| Mean ± SD (nm) | 14.4 ± 1.4 | 15.8 ± 1.4 | 15.0 ± 2.0 | 14.0 ± 2.0 | 13.7 ± 1.4 | 13.5 ± 1.9 |

**Table S2. Constructs used in this study.**

| Construct | Species | Backbone | Cav1 Sequence | Tags | Source |
|---|---|---|---|---|---|
| WT CAV1 | *Homo sapiens* | pET20 | Full-length | -LE-6His | This study |
| N173X | *Homo sapiens* | pET20 | 1-172 | -LE-6His | This study |
| V170X | *Homo sapiens* | pET20 | 1-169 | -LE-6His | This study |
| F160X | *Homo sapiens* | pET20 | 1-159 | -LE-6His | This study |
| P158P | *Homo sapiens* | pET20 | 1-158+novel C-terminus | -LE-6His | This study |
| ΔC | *Homo sapiens* | pET20 | 1-147 | -LE-6His | This study |
| CAV1β | *Homo sapiens* | pET20 | 32-178 | -LE-6His | This study |
| ΔN | *Homo sapiens* | pET20 | 1, 49-178 | -LE-6His | This study |
| ΔNΔC | *Homo sapiens* | pET20 | 1, 49-147 | -LE-6His | This study |
| MBP-CAV1 | *Homo sapiens* | pET28 | Full-length | 6His-MBP-TEV-6His-Factor Xa- | This study |
| CAV1-Venus | *Homo sapiens* | pET28 | Full-length | -TEV-Venus-10His | This study |
| CAV1β -Venus | *Homo sapiens* | pET28 | 32-178 | -TEV-Venus-10His | This study |

**Table S3. Primers used for plasmid construction in this study.**

| | |
|---|---|
| WT CAV1 | |
| Forward | 5'GCGGCCCATATGTCTGGTGGTAAATACGTTGACTCTGAAGG3' |
| Reverse | 5'GCGGCCCTCGAGGATTTCTTTCTGCAGGTTGATACG3' |
| N173X | |
| Forward | 5'GCGGCCCATATGTCTGGTGGTAAATACGTTGACTCTGAAGG3' |
| Reverse | 5'GCGGCCCTCGAGGATACGAACGTTAGAGAAGATTTTACC3' |
| V170X | |
| Forward | 5'GCGGCCCATATGTCTGGTGGTAAATACGTTGACTCTGAAGG3' |
| Reverse | 5'GCGGCCCTCGAGGTTAGAGAAGATTTTACCAACCGCTTCG3' |
| F160X | |
| Forward | 5'GCGGCCCATATGTCTGGTGGTAAATACGTTGACTCTGAAGG3' |
| Reverse | 5'GCGGCCCTCGAGCAGCGGGTCGCAAACGGTGTGAACG3' |
| P158P | |
| Forward | 5'GCGGCCCATATGTCTGGTGGTAAATACGTTGACTCTGAAGG3' |
| Reverse | 5'GCGGCCCTCGAGCTTATATTTCTTTCTGCAAGTTGATGCGG3' |
| ΔC | |
| Forward | 5'GCGGCCCATATGTCTGGTGGTAAATACGTTGACTCTGAAGG3' |
| Reverse | 5'GCGGCCCTCGAGAACACGAGAGATGCACTGG3' |
| CAV1β | |
| Forward | 5'GCGGCCCATATGGCGGACGAACTGTCTG3' |
| Reverse | 5'GCGGCCCTCGAGGATTTCTTTCTGCAGGTTGATACG3' |
| ΔN | |
| Forward | 5'GCGGCCCATATGATCGACCTGGTTAACCG3' |
| Reverse | 5'GCGGCCCTCGAGGATTTCTTTCTGCAGGTTGATACG3' |
| ΔNΔC | |
| Forward | 5'GCGGCCCATATGATCGACCTGGTTAACCG3' |
| Reverse | 5'GCGGCCCTCGAGAACACGAGAGATGCACTGG3' |
| MBP-CAV1 | |
| Forward | 5'CGGGATCCCATCATCATCATCATCATATCGAAGGTCGTATGTCTGGGGG CAAATACGTAGAC3' |
| Reverse | 5'CGGAATTCTTATATTTCTTTCTGCAAGTTGATGCG3' |
| CAV1β -Venus | |
| Forward | 5'GATATACCATGGCGGACGAACTGTCTG3' |
| Reverse | 5'CACGCCTCGAGATCTAAGCTTTCCGGAGCCCTG3' |

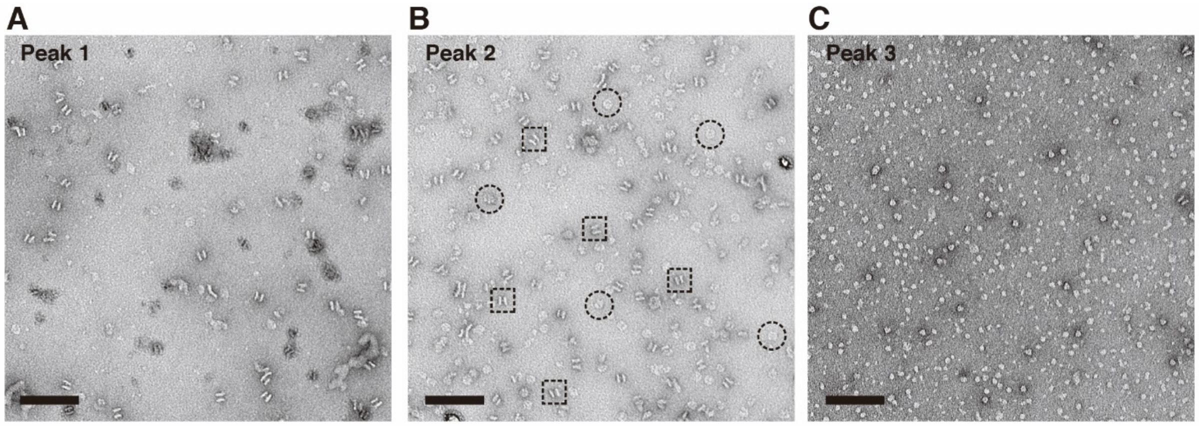


Figure S1

**Figure S1. Characterization of CAV1 complexes purified from *E. coli*.**
Representative negative stain images of particles found in Peaks 1 (**A**), 2 (**B**), and 3 (**C**) of gel filtration profile of Cav1-His6 purification. Peak 2 was used for further structural characterization. Examples of *en face* views are circled, and boxes mark examples of side views of dimerized 8S complexes. Scale bar, 100 nm.

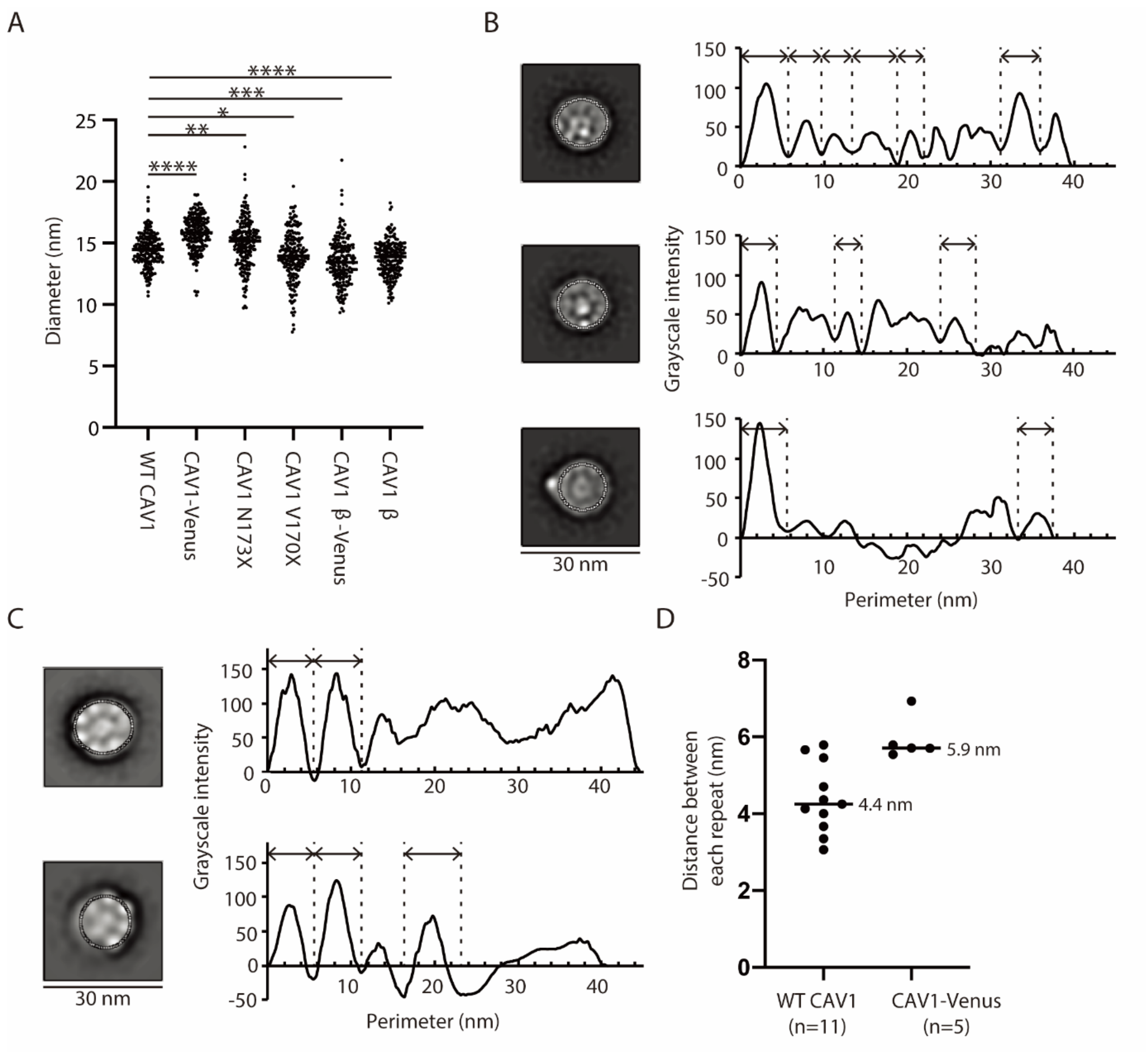


Figure S2

**Figure S2. Size measurements of negatively stained CAV1 complexes.** (**A**) Plots of diameters of 200 individual 8S complexes of WT CAV1, CAV1-Venus, N173X, V170X, CAV1β-Venus, and Cav1β seen in negative stain images. An ordinary one-way Analysis of Variance (ANOVA) test was performed to determine whether the difference in the means were statistically significant, whereas Tukey's multiple comparisons test was performed for pairwise comparisons. ****$P \leq 0.0001$; ***$P \leq 0.001$; **$P \leq 0.01$; *$P \leq 0.05$. (**B**) Oval plot profiles for three additional WT CAV1 *en face* class averages. Positions of the oval used for analysis are shown in white dotted lines on the particle images. On the graphs, dotted lines mark the boundaries between adjacent globular domains, which we interpret to represent CAV1 monomers. Note that only globular domains with distinct boundaries between them were included in the analysis. (**C**) Oval plot profiles for two CAV1-Venus *en face* class averages. (**D**) Plots of average distance each globular domain occupies for WT CAV1 and CAV1-Venus, as measured in panels B and C. n indicates the number of monomers measured.

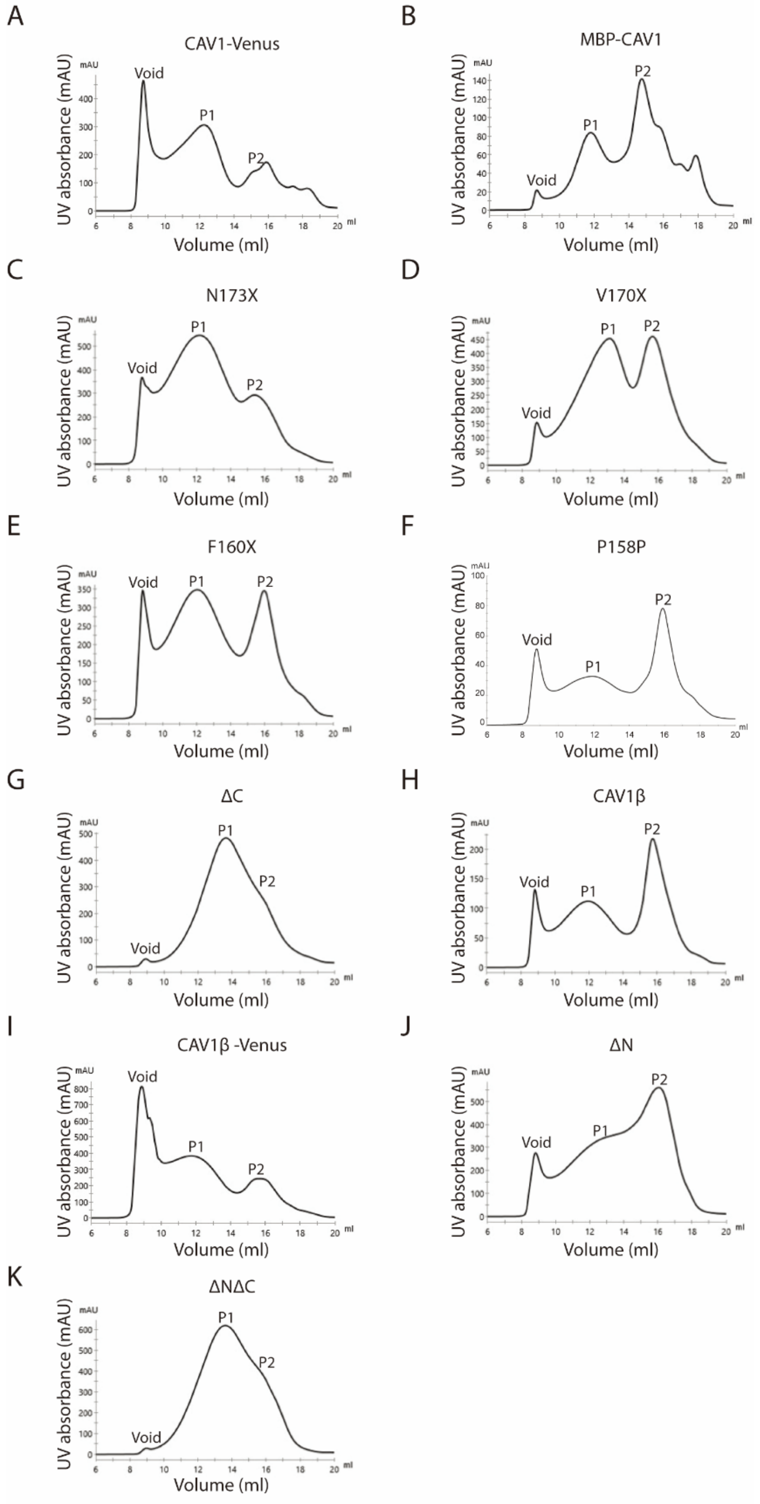


Figure S3

**Figure S3. FPLC traces of CAV1 constructs.** The indicated proteins were purified from E. coli membranes and applied to a Superose®6 Increase 10/300 GL column. Shown are the elution profiles for CAV1-Venus (**A**), MBP-CAV1 (**B**), N173X (**C**), V170X (**D**), F160X (**E**), P158P (**F**), ΔC (**G**), CAV1β (**H**), CAV1β-Venus (**I**), ΔN (**J**), and ΔNΔC (**K**). For all of the traces listed above, Peak 1 contains the fully organized oligomers and was used for structural characterization.

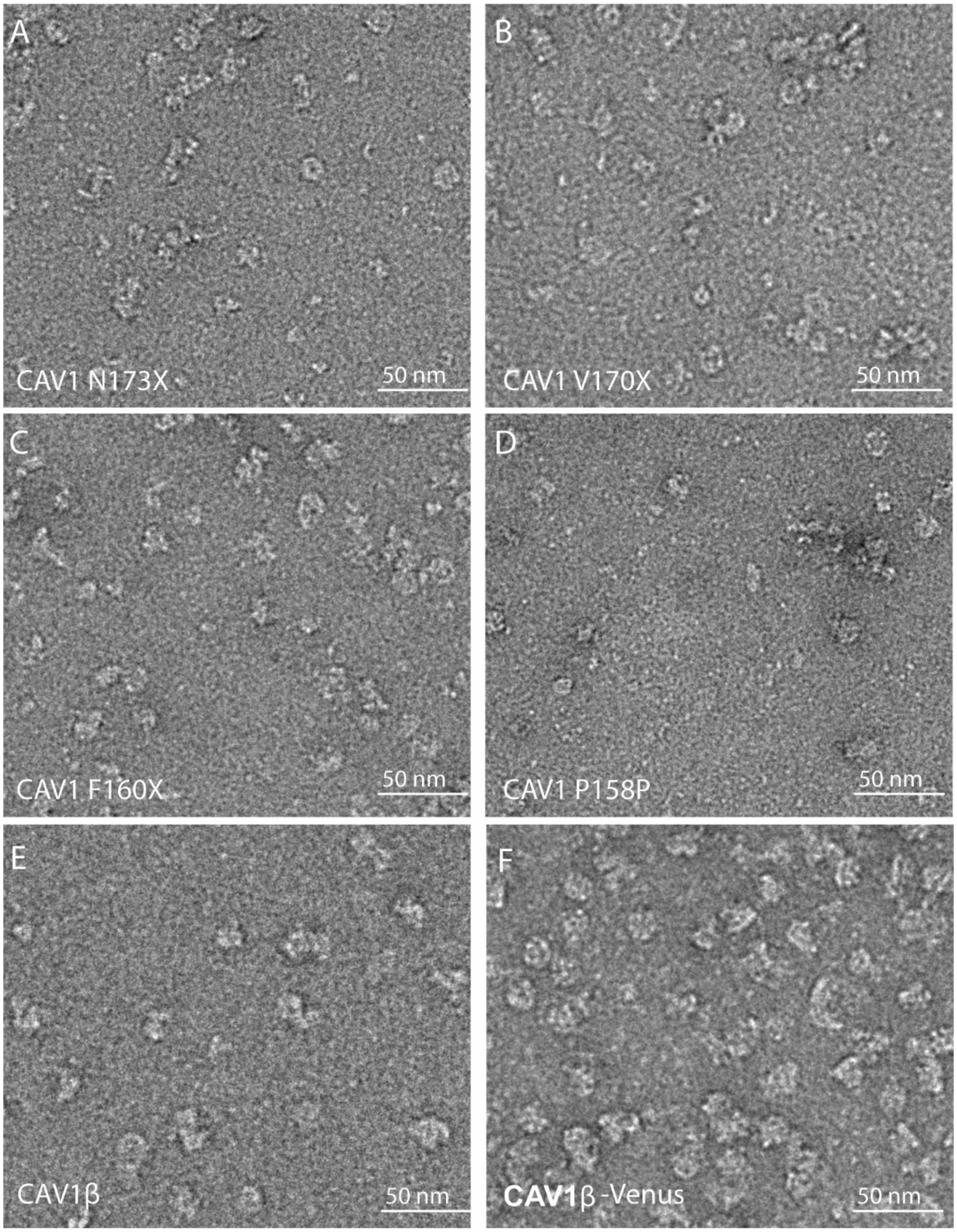


Figure S4

**Figure S4. Negative stain analysis of CAV1 C-terminus truncations and CAV1β.** Representative negative stain images of (**A**) N173X, (**B**) V170X, (**C**) F160X, (**D**) P158P, (**E**) Cav1β, and (**F**) Cav1β-Venus. Scale bars, 50 nm.
